# Peritoneal autoantibody landscape in endometriosis

**DOI:** 10.1101/2022.05.27.493373

**Authors:** Sarah Harden, Tse Yeun Tan, Chee Wai Ku, Jieliang Zhou, Qingfeng Chen, Jerry Kok Yen Chan, Jan Brosens, Yie Hou Lee

**Affiliations:** Critical Analytics for Manufacturing Precision Medicine, Singapore-MIT Alliance for Research and Technology, Singapore, Singapore; Division of Biomedical Sciences, Clinical Science Research Laboratories, Warwick Medical School, University of Warwick, Coventry CV4 7AL, UK; Institute of Molecular and Cell Biology, Agency for Science, Technology and Research, Singapore, Singapore; Department of Reproductive Medicine, KKH, Singapore, Singapore; OBGYN-Academic Clinical Program, Duke-NUS Medical School, Singapore, Singapore; KK Research Centre, KK Women’s and Children’s Hospital, Singapore, Singapore; Tommy’s National Centre for Miscarriage Research, University Hospitals Coventry & Warwickshire, Coventry CV2 2DX, UK; Centre for Early Life, Warwick Medical School, University of Warwick, Coventry CV4 7AL, UK

**Author notes:** ***Address correspondence to:*** Lee Yie Hou Ph.D., OBGYN-ACP, Singhealth-Duke-NUS Academic Medicine, 100 Bukit Timah Road, Singapore 229899, Singapore-MIT Alliance for Research and Technology, 1 CREATE Way, #04-13/14 Enterprise Wing, Singapore 138602, **E-mail:**.

**Keywords:** Endometriosis, Autoantibodies, Autoimmunity, Precision medicine, Peritoneal

## Abstract

Women with endometriosis have a profound association with autoimmunity. An excess of autoantigens in the peritoneal cavity resulting from retrograde menstruation could lead to inflammation and pathologic autoimmunity. Using a native-conformation protein array, proteome-wide analysis of autoantibodies (AAbs) against 1623 proteins were profiled in peritoneal fluids (PF) of 25 women with endometriosis and 25 endometriosis-negative women. 46% of endometriotic women have five or more AAbs. Diverse cognate autoantigens were identified and corresponding AAbs against proteins involved in implantation, B-cell activation/development, and aberrant migration and mitogenicity. AAbs recognizing tumour suppressor protein p53 were the most frequent at 35% and were targeted against native and citrullinated p53 forms. Further, unsupervised hierarchical clustering and integrative pathway analysis, we observed clusters of endometriosis-associated infertile women with 60% positive for two or more AAbs which are involved in PDGF, TGF-β, RAC1/PAK1/p38/MMP2 signaling, LAT2/NTAL/LAB-mediated calcium mobilisation and integrin-mediated cell adhesion. Together, our data identifies peritoneal autoimmunity in a significant subset of women with endometriosis, with diverse impact on infertility and disease pathophysiology.

## INTRODUCTION

Endometriosis is a common gynaecological disorder that affects up to 10% of women and is associated with chronic pain and infertility. The disease is defined by the presence of endometrial-like implants found outside the uterus, most commonly on the peritoneum ^1^. Given the severe and chronic symptoms, women often develop social and psychological morbidities ^2^, with the cost of disease estimated at US$70 billion in the United States, similar to those incurred by Rheumatoid Arthritis and Diabetes Mellitus ^3^. The estimated annual economic loss is US$22 billion due to productivity loss ^4^. The retrograde flow of shed endometrial tissues implanting onto extra-uterine sites and the harbinger of foreign endometrial tissues and cells to extra-uterine locations is commonly accepted as the principal initiator of the disease ^5^. The appearance and ineffective clearance of these foreign and cell debris including antigens in extra-uterine locations during retrograde menstruation, potentially provoke autoimmune responses, immunologic tolerance, or rejection of the autograft with alloantigenic potential ^6^.

The plausibility of endometriosis being considered an autoimmune disease has been postulated, insofar that endometriosis meets most of the classification criteria of an autoimmune disease, and there is deregulation of the apoptotic process ^7^. Since endometriotic lesions originate from autologous cells containing self-antigens, it can be speculated that it is abnormal exposure or presentation of these antigens that facilitates an autoimmune response. This follows the discovery of IgG, IgM, and IgA AAbs directed against cell-derived antigens such as phospholipids and histones ^8^. Anti-endometrial and anti-ovarian autoantibodies (AAbs) against transferrin and alpha 2-HS glycoprotein were found in PF of women with endometriosis ^9,10^. AAbs against endometrial and ovarian tissue in sera, vaginal, and cervical secretions in women with endometriosis suggest autoimmune dysregulation in women with endometriosis ^9^, and organ-specificity ^11^. Further, abnormalities in endometrial AAbs strongly suggest a role in endometriosis-associated infertility (EAI) ^12^. The association of endometriosis with autoimmune diseases and increased incidence of AAbs with endometriosis provides further support ^13–17^. Interestingly, treatment with Danazol or GnRH analogues, which are commonly used as first- or second-line therapies for the treatment of endometriosis, suppressed AAb levels ^18,19^.

A defective peritoneal environment characterizes endometriosis, by which the peritoneal fluid (PF) that bathes the peritoneal cavity and surrounds endometriotic implants is an important microenvironment that contributes to endometriosis. The PF is rife with cytokines such as IL-1□, IL-6, IL-8, IL-10, etc ^20^, and cytokines are critical for autoimmune pathophysiology and production. B-cell activating factor (BAFF or BLyS), a cytokine necessary for normal B cell development, was up-regulated in endometriosis ^21^. Pathological analyses reported the presence of plasma cells (precursors of B-cells), atypical B-cells and activated macrophages in endometriotic lesions ^21^. Intrinsic defects in peritoneal macrophages in endometriosis may also contribute to autoimmunity. Macrophages are important immune cells that maintain immune homeostasis via phagocytosis of foreign matter, apoptotic or necrotic cells, and are recruited to the peritoneum where they are prominently associated with endometriosis ^22–24^. Dysregulation in these immune cells promotes skewed tolerogenic peritoneal environments in endometriosis.

The tumour suppressor p53, encoded by the *TP53* gene, is a DNA motif binding transcription factor that governs core cellular programs to ensure cell and tissue homeostasis, including arresting cell cycle progression and apoptotic response to cellular stress ^25^. Different patterns of p53 mutations have been reported in endometriosis, including missense mutations, overexpression, deletion of the *TP53* locus, and loss of heterozygosity, that can contribute to or trigger an immune reaction by causing self-immunization of non-wildtype p53 ^26–30^. There is a strong correlation between the frequency of anti-p53 AAbs and the frequency of p53 mutations in certain types of cancer, suggesting that p53 mutations are associated with the generation of these AAbs ^26^. Studies in mice revealed the close association of p53 deficiency with the development of autoimmune and inflammatory diseases ^31,32^. In particular, monocytes/macrophages deficient in p53 inefficiently clear apoptotic and necrotic cells, and the failure to clear dying cells can lead to accumulation of autoantigens that promote further generation of autoimmunity and chronic inflammation ^31,33^.

Although examples of autoimmune responses have previously been described, the comprehensive breadth of AAb reactivities in endometriosis remains undetermined. In this study, an integrated proteome-wide and bioinformatic analysis of more than 1600 functional IgG AAbs was performed in PFs of patients with endometriosis. We found that in a subset of patients with endometriosis there is diverse autoreactivity and elevated AAbs that target biological processes related to fertility, autoimmunity, and endometriosis pathophysiology. In these patients, p53 was identified as the most frequent PF AAb target, which was present in both the native and citrullinated form. Citrullination is a post-translational modification of arginine side chains into citrulline that produces non-self-neoepitopes, dramatically altering immunogenicity and driving further autoantibody production ^34^. Stratification by p53 AAb positive patients found a monocyte / macrophage PF signature. Together, these findings have important implications for stratification in endometriosis and the development of new therapeutic strategies against a subset of patients with endometriosis.

## METHODS AND MATERIALS

### Study design and Patient Enrolment

The objective of this study was to identify AAbs found in the peritoneal fluid in women with endometriosis. Patients who underwent laparoscopic procedures at the KK Women’s and Children’s Hospital, Singapore for various indications such as suspected endometriosis, infertility, sterilization procedures and/or pelvic pain were recruited into the study. Women provided written informed consent for the collection of samples under Centralised Institutional Research Board approval (CIRB 2010-167-D).

Exclusion criteria include menstruating patients, postmenopausal patients, anovulatory patients, patients on any form of hormonal therapy for at least three months prior to laparoscopy, and other potentially confounding diseases, including diabetes, rheumatoid arthritis, inflammatory bowel disease, multiple sclerosis, and systemic sclerosis. Diagnostic laparoscopy was performed on all patients, with careful inspection of the uterus, fallopian tubes, ovaries, pouch of Douglas, and the pelvic peritoneum by gynaecologists subspecializing in reproductive endocrinology and infertility. PF was prepared as previously described ^11^, in line with the Endometriosis Phenome and Biobanking Harmonisation Project Standard Operating Procedures ^35^. The presence of endometriosis was systematically recorded and scored according to the revised American Fertility Society classification (rAFS) of endometriosis ^36,37^ and classified as women with endometriosis (EM+; *N*=25) or without endometriosis (EM-; *N*=25). Infertile women with endometriosis (EM+ EAI; *N*=15) and without endometriosis (EM-EAI; *N*=13) were extracted for the subsequent analysis. Patient characteristics are shown in **Table 1**.

**Table 1.**
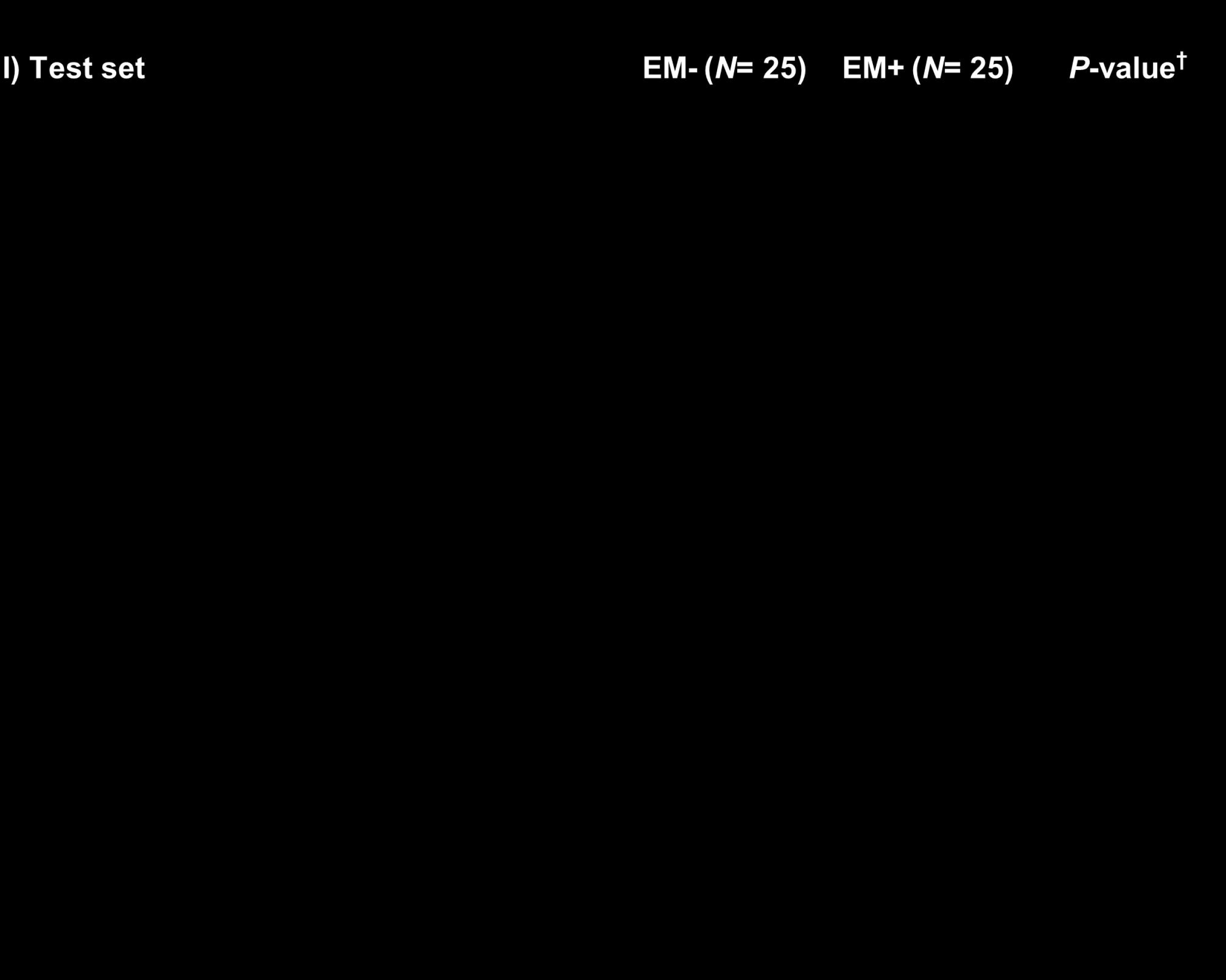
Summary of patient characteristics.

### Autoantibody proteomics

Proteome-wide AAb profiling utilised the functional protein Immunome™ microarray platform (Sengenics, UK), covering 1623 wild-type proteins (antigens), as noted in **Table S1**. This AAb protein array uses a compact, folded, biotinylated, ∼80 amino acid residue domain derived from the *E. coli* biotin carboxyl carrier protein (BCCP) that preserves the structure and function of the embedded antigens, thus offering exquisite selectivity and specificity of bounded autoantibodies ^38,39^. Patient or control PF samples were diluted 1:200 in 2 mL dilution buffer (0.1% Triton X100 (v/v), 0.1% BSA (w/v) in PBS) and applied to the array. The arrays were incubated in Quadriperm dishes (Greiner BioOne, Stonehouse, UK) and placed on a horizontal shaker at 50 rpm for 2 h at room temperature. After several washes, anti-human IgG was diluted 1:1000 in assay buffer and Cy3-rabbit antihuman IgG (Dako Cytomation) by incubation for 2 h at room temperature according to the manufacturer’
ss recommendations. The array was washed again and dried by centrifugation. All arrays were scanned at 10 µm resolution using a microarray scanner (Axon 4200AL with GenePix Pro Software, Molecular Devices, Sunnyvale, CA, USA) and fluorescence of labelled IgG was detected according to the manufacturer’s instructions. The images were saved as 16-bit tiff files and the analysis was performed using GenePix software. The interaction between microarray antigens and PF AAbs was detected as fluorescence of the bound fluorescently-labeled IgG at the protein-specific position on the microarray. The intensity of fluorescence is proportional to the amount of AAb present in the PF. Local background obtained from the control spots on the array was automatically subtracted, and relative fluorescence units (RFU) for each microarray spot were recorded. Each antigen was immobilised in quadruplicate on the array. The median RFU for the four readings of each antigen was used for further analysis. All arrays passed quality control tests. Mean of CV% of all protein replica spots across all samples was 8.12% (**Fig S1A**). Cy3-labelled biotinylated BSA (Cy3-BSA), with concentrations, kept constant across the arrays, serves as the positive control across slides and as a housekeeping probe for normalisation of signal intensities across samples, has a CV% of 8.08% (**Fig S1B**).

### Citrullination proteomic analysis

PF samples were diluted at 1:100 in dilution buffer as above. When completely thawed, each sample was vigorously vortexed three times at full speed and spun down for 3 min at 13,000 rpm using a microcentrifuge. 30 μL of the sample was pipetted into 3 mL of wash buffer containing 0.2% v/v tween-20 in 1 × PBS (20°C) and vortexed to mix three times. Human PAD1, 2, and 6 were chosen for further characterisation due to their expression in the uterus and the ovaries ^40–42^, and are therefore most relevant to endometriosis. PAD1, PAD2, PAD6 were incubated on the protein array for the enzymatic conversion of arginine to citrulline groups (herein referred to as citrullination protein array). Briefly, each slide was rinsed in 3 mL Wash Buffer for 5 min. When the slides were rinsed completely, they were blocked in CT100plus blocking buffer for 1 h. All slides were then washed three times in wash buffer for 5 min at room temperature. The slides were then incubated with 3 mL of 1 μg/mL human PAD1, PAD2 or PAD6, covered with aluminium foil and incubated for 3 h at 37°C at 50 rpm. The slides were then washed for 3 × 5 min at room temperature in 3 mL of wash buffer. The citrullinated array was then incubated with diluted PF samples on a horizontal incubator at 20°C for 2 h. The citrullination protein array was then incubated with an anti-citrulline antibody and fluorescently-labelled detection antibody to detect citrulline groups. IgG binding was detected by incubation with Cy3-rabbit anti-human IgG (Dako Cytomation) labeled according to the manufacturer’s recommended protocols (GE Healthcare). Arrays were immersed in a hybridization solution containing a mixture of Cy3-rabbit antihuman IgG solution diluted 1:1000 in wash buffer for 2 h at 50 rpm at 20°C. After incubation, the slide was washed with wash buffer, 3 × 5 min at 50 rpm at room temperature and a final wash with 1× PBS for 5 min. The slides were then dried for 4 min at 400 g at room temperature and scanned. Hybridization signals were measured with a microarray laser scanner (Agilent Scanner) at 10 μm resolution. Fluorescence levels were detected according to the manufacturer’s instructions, whereby each spot is plotted using Agilent Feature Extraction software. All samples passed QC parameters that evaluate quantitative metrics related to array and assay quality and consistency of results. The coefficient of variance (CV%) of the intra-protein, intra-slide and inter-array for all proteins and control probes of PAD1, PAD2, PAD6 citrullinated protein arrays were 9.20%, 8.35% and 6.63% respectively, below the QC limit of 15%.

### Multiplex immunoassay analysis

The levels of 48 cytokines were measured in the PF fluid using a multiplex suspension bead immunoassay (BioRad, CA, USA; **Table S1**) as previously described ^20^. Briefly, 10 μL of PF was mixed with 10 μL of primary antibody-conjugated magnetic beads on a 96 DropArray plate (Curiox Biosystems, Singapore) and rotated at 450 rpm on a plate shaker for 120 min at 25°C while protected from light. Subsequently, the plate was washed three times with wash buffer on the LT210 Washing Station (Curiox) before adding 5 μL of the secondary antibody and rotating at 450 rpm for 30 min at 25°C protected from light. The plate was washed three times with wash buffer and 10 μL of streptavidin-phycoerythrin was added and rotated at 450 rpm for 30 min at 25°C protected from light. The plate was washed three times with wash buffer. 60 μL of reading buffer was then added and transferred to a 96 conical well microtiter plate, and the samples were read using the Bio-Plex Luminex 200 (BioRad). The beads are classified by the red classification laser (635 nm) into its distinct sets, while a green reporter laser (532 nm) excites the phycoerythrin, a fluorescent reporter tag bound to the detection antibody. All samples were run in duplicates and the average was reported. Quantitation of the 48 cytokines in each sample was then determined by extrapolation to a six- or seven-point standard curve using five-parameter logistic regression modelling. Assay CV averaged <12 %. When samples were detected in less than 50% of patients or below the lower limit of quantification, they were considered undetected. Calibrations and validations were performed before runs and every month respectively.

### Data analysis

For statistical analyses, parametric t-test, and Pearson linear regression analysis were used, and statistical significance was set at *p*<0.05. Penetrance-based fold change (pFC) analysis method was implemented for the identification of highly expressed proteins in each case sample. This method removes any false positive signals from the data by setting a protein-specific threshold (i.e. background threshold). This defined per-protein background threshold is calculated on the basis of two standard deviations from the mean signal intensities for each specific antigen measured of the control group. Any antigens below this threshold are excluded from further analysis. Individual fold changes for both case and control are calculated by dividing the RFU value for each protein in each sample, H, by the mean of the RFU values of each protein across all the control samples (i.e. background threshold), that is:

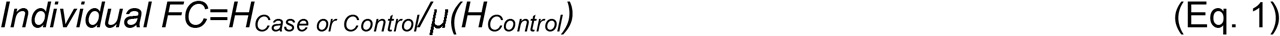

For proteins with an individual fold change less than 2 fold above the background threshold, their signal intensities (RFU) are replaced with zeros. Penetrance frequency (number of case and control samples with individual fold changes ≥ 2-fold) for both case and control were determined for each protein as follows:

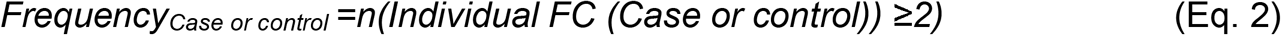

Penetrance Fold Changes (pFC) for both the case and control groups are calculated for each protein as follows:

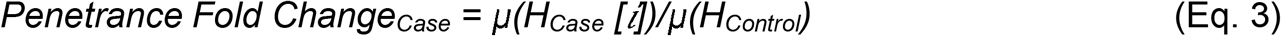

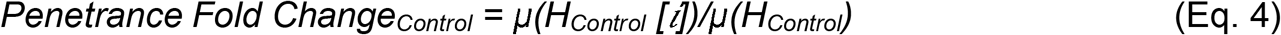

*Where H*_*Case*_ *[i]= H*_*Case*_ *with Fold Change*_*Case*_ ≥*2 fold* and *H*_*Control*_ *[i]= H*_*Control*_ *with Fold Change*_*Control*_ ≥*2 fold*

Putative endometriosis-associated AAbs were identified and ranked according to the following criteria:

1. Penetrance Fold Change_Case_ ≥ 2, and
2. % Frequency_Case_ ≥ 20%

### Pathway enrichment bioinformatics analysis

Functional enrichment data was obtained from ToppGene Suite based on Gene Ontology, pathways and disease, using the ToppFun tool ^43^. *P*-values were calculated using the probability density function and false discovery rate (FDR) Benjamini and Hochberg corrected, with a corrected p-value or q-value cutoff of *q* = 0.05. The OpenTargets Platform was utilised to find proteins associated with autoimmunity and endometriosis ^44^. Here, Overall Association scores between a target and disease were calculated using data from multiple sources, and adjusted depending on data source and data type. A value of 1 represents the most associated. These proteins were cross-referenced to the proteins identified as autoantibodies in the Immunome™ microarrays, to highlight proteins known to be involved in autoimmuinity, endometriosis, or both.

## RESULTS

### Diversity of peritoneal autoantibodies in a subset of endometriosis patients

To obtain a global overview of the repertoire of immunoreactive IgG AAbs in endometriosis, PF samples from 25 EM+ women and 25 EM-controls (**Table S2**) were analysed for functional human AAb levels against 1623 correctly folded full-length antigenic proteins. In EM+, 84.1% of PF AAbs were found at frequencies of four to six (**Figure 1A**). This contrasts with EM-controls whereby the majority of PF AAbs (95.1%) were presented with a low frequency of one and two (Kruskal Wallis test corrected with Dunn’s test for multiple comparisons, *p*<0.0001). Of the analysed 1623 IgG AAbs, 351 discrete AAbs were putatively considered as significant (pFC_Case_ ≥ 2, penetrance Frequency_Case_ ≥ 20%; **Table S3**). Previously identified PF and endometrial tissue AAbs in endometriosis, anti-histone H1.2 and anti-alpha 2-HS-glycoprotein (AHSG) AAbs were validated (pFC = 3.59 and 3.24; penetrance Frequency_Case_ = 20% and 24%, respectively; **Figure S2A, B**) ^12,45,46^. By hierarchical clustering, a cluster of twelve (46%) EM+ cases with strong autoimmune profiles (five or more significant AAbs per patient; average pFC = 2.24 versus average pFC = 0.67 for the rest of EM+) were observed (**Figure 1B, C**), although the cluster was not associated with ASRM endometriosis severity, menstrual phase, age, or fertility. Intriguingly, a cluster of five fertile EM-controls displayed mild AAb positivity (average pFC =1.5 compared to average pFC = 0.71 for the rest of EM-). Many of the EM+ AAbs include markers of fertility such as decidualization (PRL) ^47^ and implantation (ACVR2A, SMAD5) ^48^, autoimmunity such as B-cell activation and development (BANK1, FLI1) ^49,50^, and endometriosis pathophysiology such as migration (TIMP3, MMP24) ^51,52^, and mitogenicity (PDGFB, PDGFRL, FGFR1, FGFR2, IGF2, VEGF-D) (**Figure 1B**). Pathway enrichment, incorporating data from KEGG, Reactome and BIOCARTA, indicated that the AAbs were found to elicit pathways involved in MAPK (*q* = 4.68 × 10^−7^), PDGF (*q* = 5.51 × 10^−7^), LKB1 (*q* = 2.74 × 10^−6^), FGF (*q* = 4.81 × 10^−6^), and IL-2-mediated signaling (*q* = 3.05 × 10^−6^), as well as the Toll-receptor signaling cascade (*q* = 1.13 × 10^−6^) (**Figure 1D, Table S4**). The 351 AAbs were cross-referenced to potential targets under “Autoimmunity” and “Endometriosis” disease categories using the OpenTargets Platform. 149 AAbs were found to be associated with “Autoimmunity”, four with “Endometriosis” and 74 AAbs overlapped with both “Endometriosis” and “Autoimmunity” (**Figure 1E**; **Tables S5, S6**) (50).

**Figure 1.**
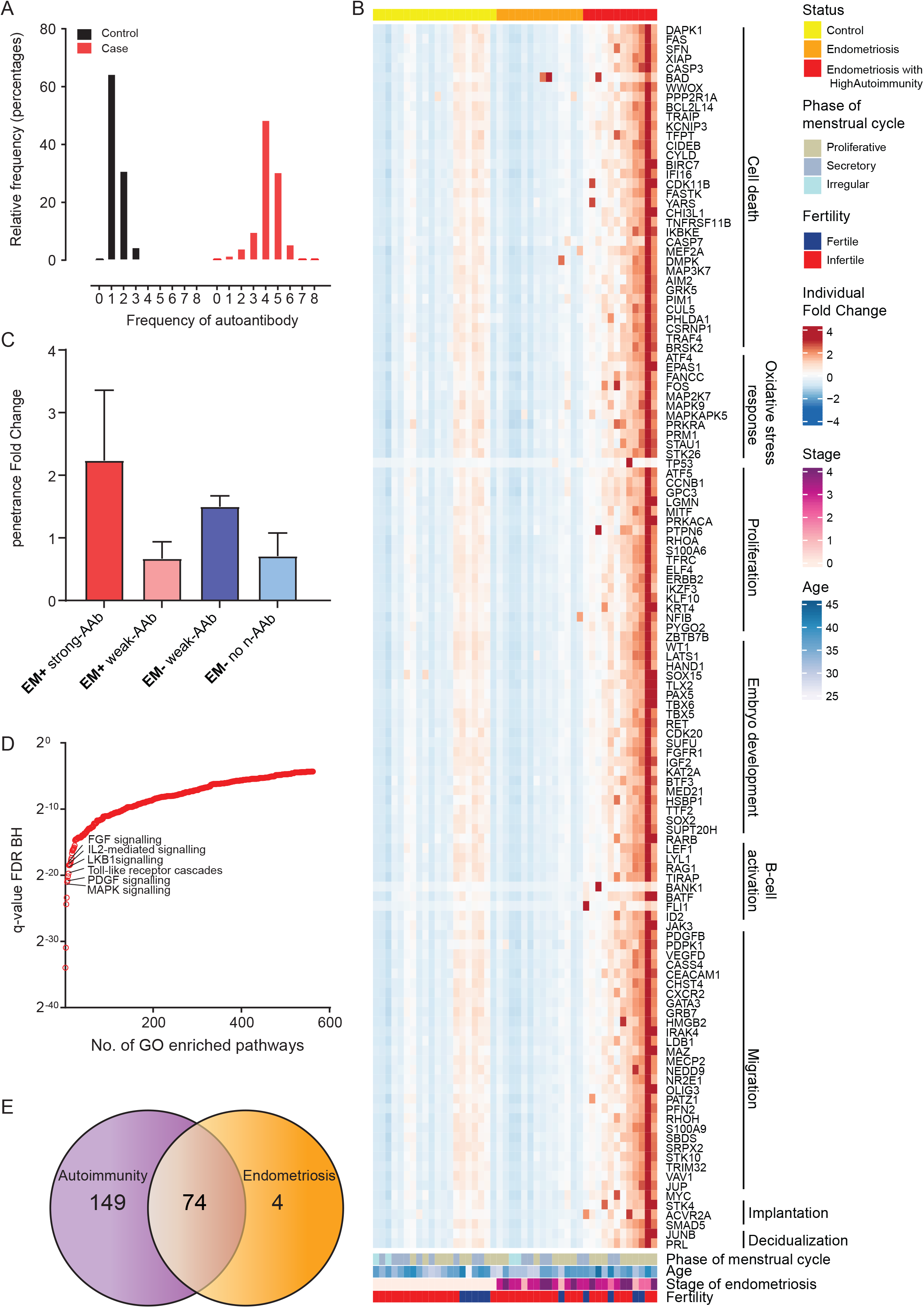
Women with endometriosis have diverse peritoneal autoantibodies. (A) Distribution histogram of frequency of positive autoantibodies with fold change ≥2, showing that more EM+ cases (n=25) possessing frequencies of autoantibodies of four and five and up to eight, compared with EM-controls of one or two autoantibodies (n=25). (B) Heatmap of 351 autoantibody levels in women with (n = 25) and without endometriosis (n =25). Autoantibody levels were Z score normalised against control population mean and SD, with Z scores >2 corresponding to positive autoantibody levels. Autoantibodies cluster into three major clusters, control, endometriosis, and endometriosis with high autoimmunity. (C) Bar graphs of EM+ cases and EM-controls with strong (≥5 significant autoantibodies per patient) and weak (<5 significant autoantibodies per patient) autoimmune profiles. (D) Pathway enrichment q-values (FDR Benjamini-Hochberg), indicated that positive autoantibodies in endometriosis were found to elicit pathways involved in MAPK (*q* = 4.68 10^−7^), PDGF (*q* = 5.51 10^−7^), Toll-receptor signaling cascade (*q* = 1.13 10^−6^), LKB1 (*q* = 2.74 10^−6^), IL-2-mediated signaling (*q* = 3.05 10^−6), and^ FGF (*q* = 4.81 10^−6^)as top ranked pathways. (E) Number of the 351 autoantibodies associated with autoimmunity, endometriosis or bose, as determined by the OpenTargets Platform.

Endometriosis is associated with an aberrant cytokine and complement response (13,35). We did not find evidence of autoimmunity against notable cytokines or chemokines (not detected or low autoreactivity with canonical and non-canonical cytokines or receptors IL1A, IL13, IL18, IL37, CXCR2, CXCR4, CXCR6; **Figure S2C-J**), arguing that they are not targeted directly or neutralized by PF AAbs in endometriosis. Collectively, these results revealed that a significant subset of EM+ patients are autoimmune positive, evidenced by a relatively large number of prominent peritoneal AAbs, which target a diverse range autoantigens related to fertility and endometriosis pathophysiology.

### Elevated anti-native and citrullinated p53 antibodies in endometriosis is associated with monocyte-associated cytokine profile

The most frequently occurring EM+ AAb was anti-tumour protein p53, which was found to occur in 35% of EM+ patients and 58% in the high autoimmunity EM+ cases (**Figure 2A**). Anti-p53 AAb was significantly elevated in EM+ patients compared to EM-patients (average pFC_case_= 6.46 versus average pFC_control_ = 0), and was not associated with ASRM stage, age, menstrual phase. PF of EM-patients did not show positivity for anti-p53. Therefore, anti-p53 AAb was used to stratify EM+ patients into p53^high^ (pFC_p53_ > 2.0) and p53^low^ (pFC_p53_ < 2.0) for further investigation of whether the presence of anti-p53 AAb influenced the peritoneal inflammatory environment and the frequency of citrullinated p53.

**Figure 2.**
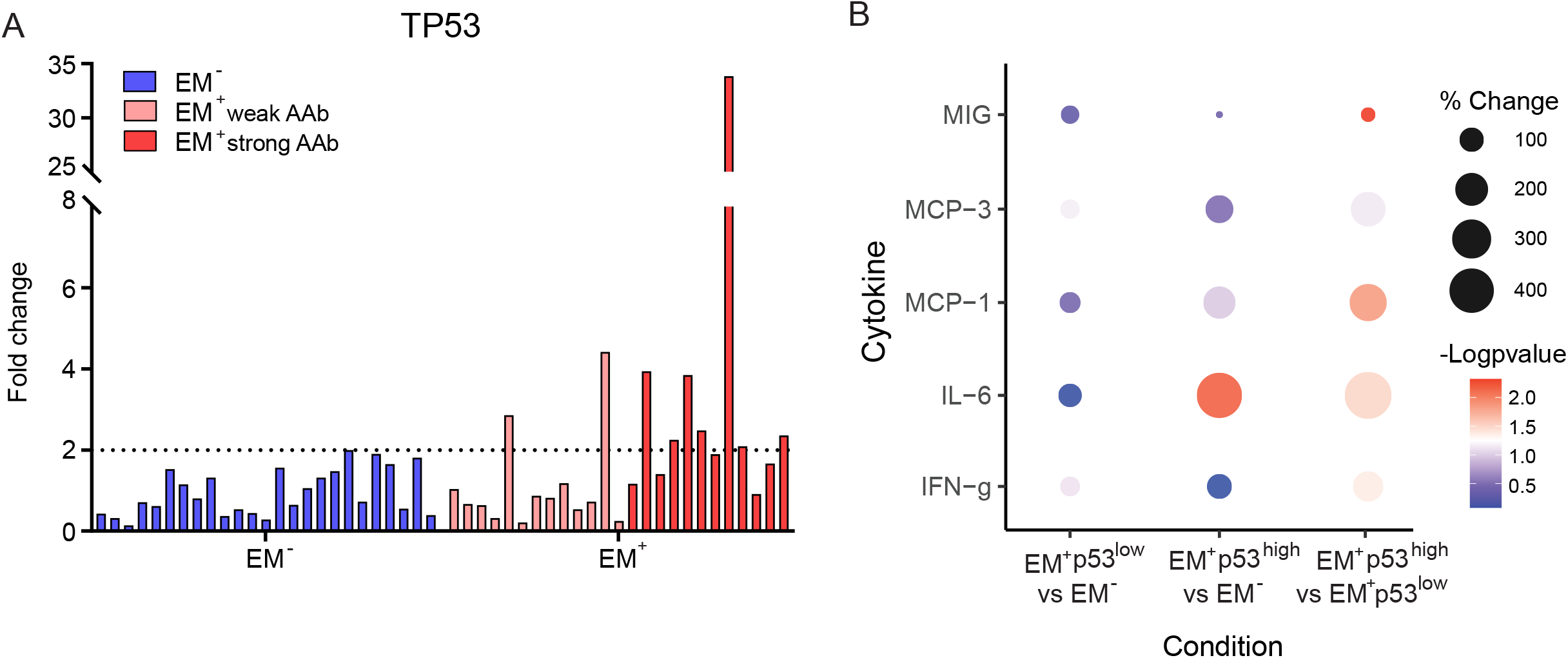
Predominance of anti-p53 autoantibody in endometriosis. (A) Bar graph of anti-p53 autoantibody fold change, showing that the autoantibody levels are elevated in EM+ cases. (B) Dot plot of chemokines indicative of changes in cytokine milieu in endometriosis patients with high p53 autoimmunity.

Because p53 provokes inflammatory responses by modulating immunological changes ^53,54^, we hypothesized that the presence of anti-p53 AAbs functionally altered the PF cytokine milieu. To test this, we performed multiplex suspension bead immunoassay of PF cytokines on EM+ p53^high^ samples compared to EM+ p53^low^ and EM-samples. 40 PF cytokines were detected (**Table S1**). A striking monocyte/macrophage-related chemokine signature comprising of significantly elevated interleukin-6 (IL-6), interferon-gamma (IFNγ), monocyte chemokine protein-1 (MCP-1), MCP-3, and reduced monokine induced by gamma interferon (MIG) distinctly marked p53^high^ samples (**Figure 2B**). When comparing p53^high^ to EM-, IL-6 levels were significantly higher in p53^high^.

Citrullination is a post-translational modification by which arginine is converted to citrulline by peptidylarginine deiminases (PADs) unveiling novel antigenic epitopes that are over-enriched in several autoimmune diseases ^55^. For the study of citrullinated targets, we pooled PF samples from EM+ patients based on their levels of anti-p53 AAb into two anti-p53 AAb groups (p53^high^ and p53^low^) (**Figure 3A**). The antigens embedded in the protein array were then citrullinated *in vitro* with PAD isoforms 1,2 and 6 and probed using anti-citrullinated antibodies in the PF samples. To validate the performance of this assay, we assessed the concordance of the generated proteome data with known citrullinated targets. Known citrullinated proteins keratins (KRT15 and KRT19), vimentin (VIM) and aldolase (ALDOA) were observed, thereby validating the assay (**Figure S3A**). The anti-p53 AAb groups had different citrullination patterns depending on whether they were incubated with PAD1, 2, or 6, with PAD1 generating the most reactive autoantigens (**Figure 3B, Figure S3B**). In the p53^high^ group 72 citrullinated AAbs overlapped in AAbs generated by the three PADs and in the p53^low^ group only one overlapped. Overall, the list of anti-citrullinated AAbs overlapped minimally with that of non-citrullinated AAbs. Interesting, citrullinated p53 was the only target among the anti-citrullinated AAbs that were common to PAD1, 2 and 6 and that overlapped with non-citrullinated AAbs (**Figure 3C**). It was approximately 1.6 times higher in the p53^high^ group compared to p53^low^ EM+ and EM- (**Figure 3D**). Together, our data suggest that in a subset of EM+ patients, their anti-p53 AAbs recognized both the modified and native form of the antigen, and elicited a unique monocyte/macrophage-like cytokine response.

**Figure 3.**
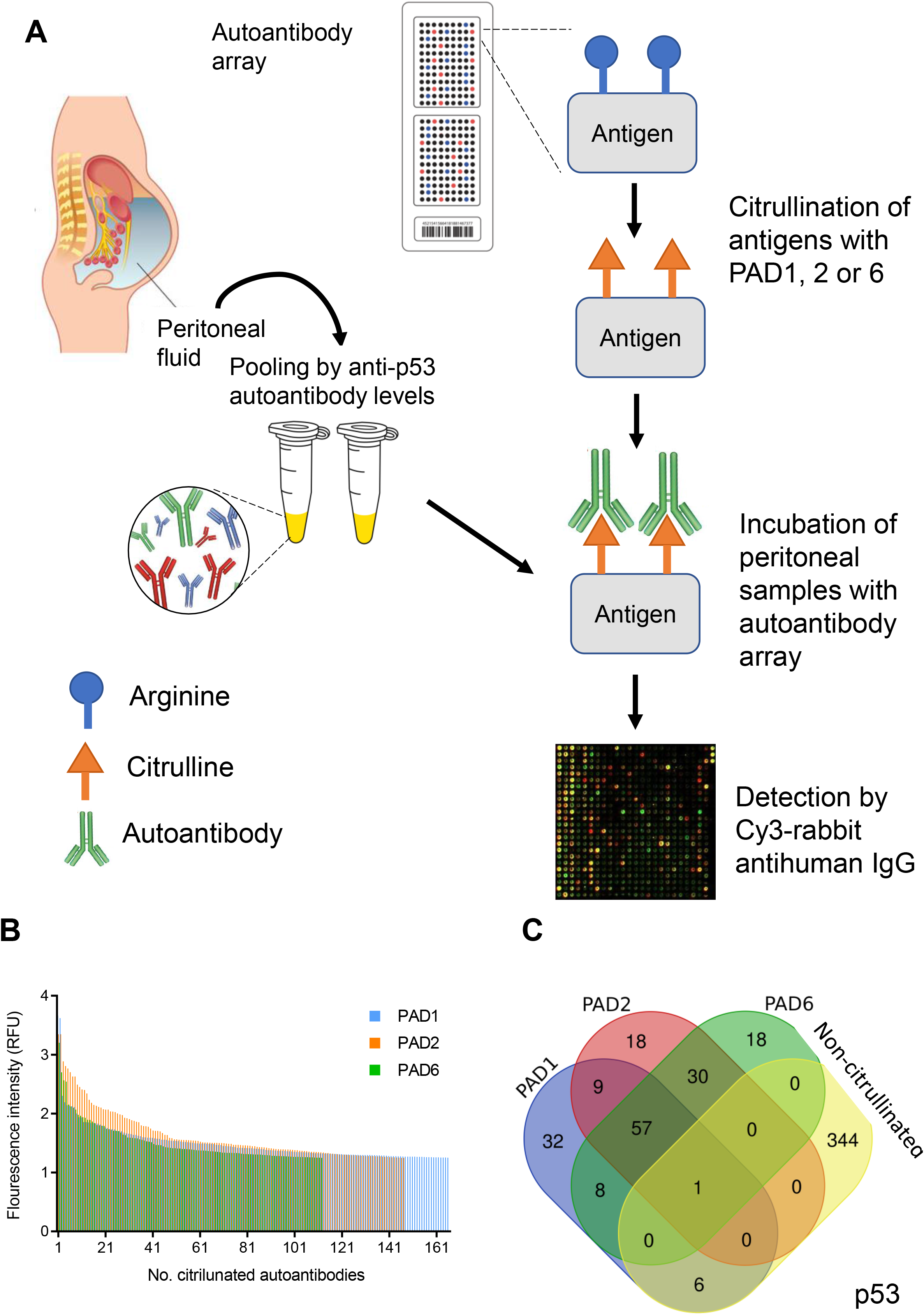
Identification of citrullinated autoantibodies in endometriosis. (A) Schematic of citrullinated autoantibody protein array generation for profiling. Peritoneal fluids from EM+ patients based on their levels of anti-p53 autoantibody levels into two anti-p53 autoantibody groups - p53^high^ and p53^low^. (B) Different citrullination patterns were obtained depending on whether they were incubated with PAD1, 2, or 6, with PAD1 generating the most reactive autoantigens. (C) In the p53^high^ group 72 citrullinated autoantibodies overlapped in autoantibodies generated by the three PADs and in the p53^low^ group only one overlapped. Overall, the list of anti-citrullinated autoantibodies overlapped minimally with that of non-citrullinated autoantibodies. Citrullinated p53 was the only target among the anti-citrullinated autoantibodies that were common to PAD1, 2 and 6 and also that overlapped with non-citrullinated autoantibodies.

### Discovery of novel peritoneal autoantibodies and p53 in endometriosis-associated infertility

Autoimmunity in EAI potentially works through different putative mechanisms or etiology ^56^. Additionally, a cluster of fertile EM-patients with a mild AAb profile was observed (**Figure 2A**) which suggested plausible confounding of fertility-associated autoimmunity in fertile individuals. This led us to perform additional analysis on 15 infertile EM+ patients and 13 infertile EM-age and ethnicity matched healthy controls selected from the above study (**Table S2**). 109 AAbs were elevated in EAI EM+ subjects (pFC≥2; Penetrance Frequency_Case_ ≥ 20%). Using Manhattan clustering distance around the median, clusters of EAI cases with positive AAb responses can be seen, with the majority (60% or 9/15) testing positive for multiple AAbs (≥2 elevated AAb) (**Figure 4A**). No AAb was elevated in infertile EM-individuals. The most prevalent AAb in EAI patients was anti-TATA-box binding protein associated factor 9 (TAF9) (33.3% frequency). Anti-p53 AAb was prevalent at 27% frequency in EAI (**Figure 4B**). 27% of them were also positive for anti-BCL2 Associated Agonist Of Cell Death (BAD), anti-fibroblast growth factor receptor 2 (FGFR2), anti-SEPTIN4, anti-nuclear nucleic acid-binding protein C1D, anti-nuclear factor erythroid 2-related factor 2 (NRF2 or NFE2L2), anti-mitogen-activated protein kinase 1 (MAPK1), anti-ETS-related transcription factor (ELF1), anti-casein kinase I isoform gamma-1 (CSNK1G1), anti-coenzyme Q8A (COQ8A), anti-ataxin-3 (ATX3) and anti-DNA-directed RNA polymerase I subunit RPA12 (ZNRD1) (**Table 2**). When examining AAbs that were common between endometriosis and autoimmunity, analysis using the OpenTargets Platform revealed 24 proteins (22%), including p53, that were found to be associated with both endometriosis and autoimmunity (**Table S7**). Integrative pathway analysis demonstrated enrichment in AAbs involved in PDGF signaling (*q* = 7.62 × 10^−5^), TGF-β signalling (*q* = 0.0007), LAT2/NTAL/LAB mediated calcium mobilisation (*q* = 0.0007), integrin-mediated cell adhesion (*q* = 0.0009), and the RAC1/PAK1/p38/MMP2 signaling axis (*q* = 0.0009) (**Table S8**).

**Figure 4.**
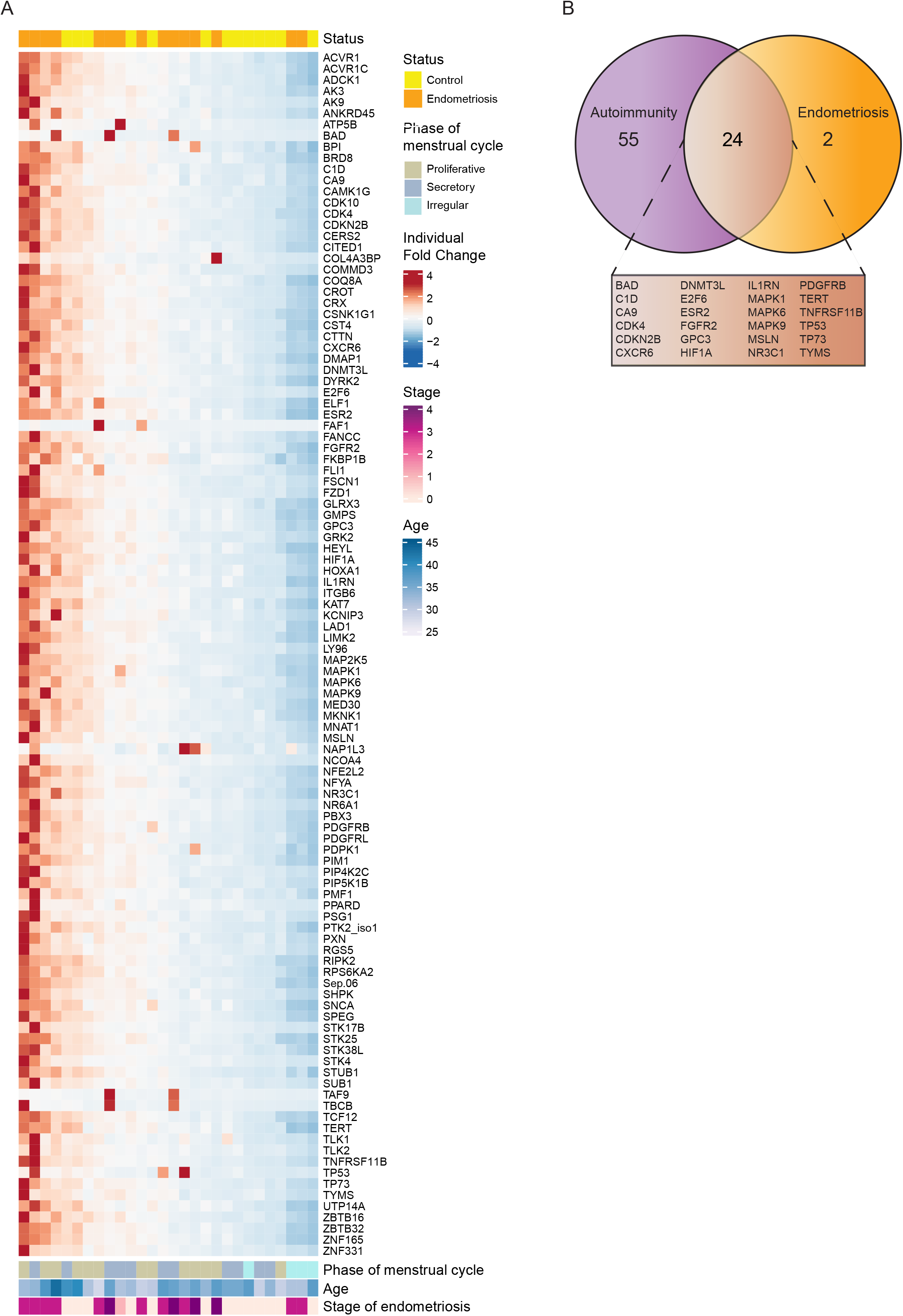
Peritoneal autoantibodies in endometriosis-associated infertility. (A) Unsupervised clustering heatmap of 109 autoantibody levels in women with (n = 15) and without endometriosis (n =13). Autoantibody levels were Z score normalised against control population mean and SD, with Z scores >2 corresponding to positive autoantibody levels. Hierarchical clustering was completed using the median distance, and Manhattan clustering. (B) Number of the 109 autoantibodies associated with autoimmunity, endometriosis or both, as determined by the OpenTargets Platform.

## DISCUSSION

Endometriosis is epidemiologically linked to autoimmune diseases such as Graves’ disease, systematic lupus erythematosus (SLE), Sjogren’s syndrome, rheumatoid arthritis and multiple sclerosis ^7,8,13–17^. Peripheral and endometrial AAbs that are associated with endometriosis have been reported ^8,57^. In this study, we investigated 1623 immunoreactive antigens against extracellular IgG AAbs found in PF. We show that there is a diversity of AAbs in women with endometriosis, with the results obtained herein confirming earlier studies on the patient’s PF that recognize the same set of antigens, including AHSG and histones. 46% of them have five or more AAbs, consistent with the observation that a proportion of EM+ patients, and not all, have an autoimmune component ^58,59^, which has led to a mix of correlation of AAbs between the severity of endometriosis and AAbs ^60–63^.

The female preponderance to an increased likelihood of autoimmunity and endometriosis-associated autoimmunity might be explained by estrogen and retrograde menstruation. Activation-induced deaminase (AID) deaminates cytosines at immunoglobulin loci, initiating a cascade of events that lead to somatic hypermutation and class switch recombination, turning IgG AAbs pathogenic. AID has been reported to be estrogen-induced ^64^. Estrogen up-regulates AID transcript and protein levels, and in ovarian tissues where estrogen levels are high, deleterious insertions of point mutations or the resolution of double-strand breaks potentially accumulate over time, generating pathogenic AAbs ^64^. Loss or deficiencies in the removal of apoptotic cells or intracellular proteins due to their release from dying cells are known to cause autoimmunity and chronic inflammation ^65,66^. The presence of cellular debris in the peritoneal cavity as a result of retrograde menstruation and defective clearance of apoptotic cells and proteins in endometriosis ^67^ presents a favourable environment that results in abnormal exposure of autologous antigens to the immune system and therefore the initiation of an autoimmune response in the peritoneal environment.

In our study, we showed diverse autoimmune responses that varied from patient to patient with endometriosis. In autoimmune diseases, auto-reactive lymphocytes expand polyclonally, have different antigen receptors on their surface, and thereby recognise different targets. AAbs are stable over long periods ^68,69^, even in the presence of low corresponding antigen levels ^70,71^. Therefore, AAb levels might be generated at different phases of endometriosis disease development, potentially early in the course of disease development, and persisted. Notably, in autoimmune disorders, such as SLE, diabetes, autoimmune ovarian failure, myasthenia gravis, and Addison’s disease, the presence of AAbs often precedes the development of clinical symptoms and potentially change over time ^8,72^. Taken together, this may explain the observed lack of correlation between AAb levels and the stage of endometriosis in this study and the heterogeneous nature of the AAb profiles. It is also intriguing to consider plausible correlates of AAb diversity across endometriosis subphenotypes ^20^, menstrual phase and age, characteristics which did not reveal an association with specific autoantibody response, although the small sample size may limit interpretation.

B-cells, T-cells, and macrophages are key immune cells in the maintenance of immune tolerance, and conversely, their dysfunction in autoimmunity. Macrophages are important immune cells that maintain immune homeostasis via phagocytosis of foreign matter, apoptotic or necrotic cells and are recruited to the peritoneum and prominently associated with endometriosis ^22–24^. Macrophages play a central role in the homeostatic clearance of cell corpses and debris by presenting antigens to T and B cells and secreting cytokines that direct the responses of T and B cells. Failure to effectively clear these cell debris will lead to the accumulation of autoantigens and drive AAb generation. Tan *et al*. recently described five subtypes of macrophages in endometriosis using single-cell RNAseq in various tissue and physiological compartments that vary between the EM+ subphenotypes that plausibly promote a tolerogenic environment for the survival of endometriotic lesions ^73^. Previous flow cytometry, cytokine and single cell-RNA sequencing analyses demonstrated the aberrant activation and dysfunction of B cell and T cell responses in endometriosis ^20,73,74^. In EM+ PF, activated T cells and relative frequencies of CD8+ T-cells, T_reg_ cells, and T_eff_ cells are higher relative to blood ^75^. CD4+Foxp3+ T_reg_ cells are significantly increased in EM+ peritoneal fluids and lesions suggesting endometriotic tolerance ^74,76,77^. Additionally, PF CD69+ T cells, which denote activated T cells, are significantly higher in EM+ compared to EM-. In the same study, CD69 T cells are strongly associated with CD161, a marker of multiple sclerosis ^78^. Recent findings in endometriosis revealed that peritoneal T_reg_ cells are disproportionally increased and functionally altered in PF and endometriotic lesions ^74,76,77^. Intrinsic defects in macrophages in endometriosis plausibly contribute to autoimmunity. However, the presence of diverse AAbs in the PF suggest eventual surmounting of autoimmunity and the introduction of self-tolerance to cell debris and lesions.

As evidenced in Proteinatlas, p53 is barely detectable in endometrial stromal or epithelial cells ^30,79^ but is highly expressed in monocytes, B- and T-cells ^80^. Although p53 expression in endometriosis has been controversial ^81–83^, our data suggest and are consistent with the finding that anti-p53 AAb is restricted to patients with mutated forms of p53. Different patterns of TP53 mutations have been reported in endometriosis, including missense mutations and loss of heterozygosity that can contribute to or trigger an immune reaction by causing self-immunization of non-wildtype p53 ^26–28^. It is worth noting that immunogenic epitopes are mapped primarily to both the N- and C-terminals of the protein ^84^, while *TP53* mutations are concentrated in the central portion (exons 4 to 8) ^85^ where autoimmunity against p53 can be generated ^86^. Therefore, the anti-p53 AAbs captured by the array used here targeted both wild-type and mutant p53 in the PF of women with endometriosis. The interplay of p53 and macrophages has also been observed, more specifically induction of anti-p53 AAbs was associated with a monocyte/macrophage cytokine signature in EM+ women. Chemokines MCP1 (CCL2) and MCP3 (CCL7) are monocyte/macrophage chemoattractants ^87^. Their increased levels point towards the infiltration of monocytes/macrophages into the peritoneum of EM+ women. The secretion of MIG (CXCL9) by predominantly monocytes/macrophages is induced by IFNγ and mediated by the JAK-STAT signalling pathway. Further studies are required to dissect the functions of anti-p53 AAbs and others, whether they are protective or pathological, and their relative contribution to endometriosis and EAI.

Citrullinated p53 has previously been reported ^88,89^ but not in endometriosis. This study showed that women with endometriosis have native and citrullinated anti-p53 AAbs, among others. To date, five human PAD isotypes that are responsible for enzymatically modifying arginine to citrulline have been described ^55^. PAD1 is expressed mainly in the uterus, PAD2 have widespread distribution including expression in the uterus, PAD3 is found in the epidermis, PAD4 expression is mainly restricted to leukocytes and PAD6 is expressed in ovaries and is a crucial enzyme in fertility ^40,41,90,91^. The protein array used in this study adopts the correct folding of proteins ^38^, providing a more accurate reflection of *bona fide in vivo* functional citrullination. The incubation of PF with citrullinated antigens converted by PADs 1, 2, and 6 identified a diversity of anti-citrulline AAbs, including p53, suggesting the likelihood that PADs can be found in the peritoneum and peritoneal cavity are responsible for citrullinating proteins. Our finding of anti-citrullinated p53 AAbs further renders p53 as a potential target of pathologic autoimmunity in endometriosis. In-depth profiling studies on the cellular and tissue distribution and expressions of PADs in endometriosis will identify the sources of citrullination and the degree of excessive and pathological conversion of proteins by the PAD isotypes.

The relatively tight clustering of fertile non-endometriosis controls who were positive for AAbs is rather striking. Fetal-derived DNA due to fetal microchimerism has been found in maternal blood as early as the first trimester and up to 38 years after pregnancy ^92,93^, triggering autoimmune adverse responses ^94^. Such observations could be explained via the following mechanism: AAbs that recognize different surface proteins might influence different stages of the fertilization process; AAbs directed against different epitopes of the same oocyte surface antigen may also affect the development and/or fertilization process, and the use of patients with a mix of fertility status. In addition, the delayed diagnosis of endometriosis ^95^ implies that by the time endometriosis is diagnosed, her ovaries might already be damaged or follicular supply exhausted, and presumably, also the target antigen for the autoimmune attack on her ovary. Interestingly, anti-ceramide transport protein (CERT, also known as COL4A3BP) AAbs were elevated in 20% of cases compared to 6.7% of controls, consistent with our earlier sphingolipidomic analysis of aberrant sphingolipid and ceramide metabolism in EAI ^96^. The impact of pregnancy on the maternal and fetal systemic immune systems can also modulate ongoing autoimmune diseases and trigger their development ^97^. The enrichment of NTAL/LAB/LAT2 pathway, found in activated B cells and monocytes ^98^, further suggests the implication of B-cell mediated autoimmunity in EAI. Integrin-mediated and TGF-β pathways have also been implicated in fertility and endometriosis ^99–101^.

In summary, this study provides an expansive peritoneal AAb landscape in a subset of endometriosis patients and identified p53 as a high-frequency AAb target that defined its association in autoimmunity. These results suggest causal inference of p53 and previously underappreciated pathways that are linked to the autoimmunological etiology of endometriosis, with implications on novel therapeutic paradigms centered around modulating these pathways and potentially immune cells to enhance endometriosis immunotherapies.

## Acknowledgements

This study was funded by Singapore Ministry of Health’s National Medical Research Council (NMRC-OFYIRG16may012 and NMRC/CG/M003/2017). National Research Foundation Singapore Fellowship (NRF-NRFF2017-03) to QC, and Wellcome Trust Investigator Award to JB (212233/Z/18/Z). Sarah Harden is supported by the University of Warwick Medical School and A^*^STAR, Singapore as part of the A^∗^STAR Research Attachment Program. CWK and JKY are supported by the National Medical Research Council, Ministry of Health, Singapore (NMRC/MOH-000596-00 and NMRC/CSA-SI-008-2016, respectively).

## Author contributions

SH and JLZ performed proteomics and cytokine analysis and analyzed data. JKYC, TTY and KCW recruited patients, provided patient samples and provided clinical feedback. YHL, SH and JJB wrote and revised the paper. YHL conceived and designed the study. YHL and JJB supervised the study.

## References

1. Saunders, P. T. K. & Horne, A. W. Endometriosis: Etiology, pathobiology, and therapeutic prospects. Cell 184, 2807–2824 (2021).

2. Siedentopf, F., Tariverdian, N., Rücke, M., Kentenich, H. & Arck, P. C. Immune Status, Psychosocial Distress and Reduced Quality of Life in Infertile Patients with Endometriosis. Am. J. Reprod. Immunol. 60, 449–461 (2008).

3. Simoens, S. et al. The burden of endometriosis: costs and quality of life of women with endometriosis and treated in referral centres. Hum. Reprod. 27, 1292–9 (2012).

4. Ahn, S. H., Singh, V. & Tayade, C. Biomarkers in endometriosis: challenges and opportunities. Fertil. Steril. 107, 523–532 (2017).

5. Wimalachandra, D. et al. Long-chain glucosylceramides crosstalk with LYN mediates endometrial cell migration. Biochim. Biophys. Acta - Mol. Cell Biol. Lipids 1863, 71–80 (2018).

6. Matarese, G., De Placido, G., Nikas, Y. & Alviggi, C. Pathogenesis of endometriosis: natural immunity dysfunction or autoimmune disease? Trends Mol. Med. 9, 223–8 (2003).

7. Nothnick, W. B. Treating endometriosis as an autoimmune disease. Fertil. Steril. 76, 223–231 (2001).

8. Eisenberg, V. H., Zolti, M. & Soriano, D. Is there an association between autoimmunity and endometriosis? Autoimmun. Rev. 11, 806–814 (2012).

9. Mathur, S. et al. Autoimmunity to endometrium and ovary in endometriosis. Clin. Exp. Immunol. 50, 259–66 (1982).

10. Mathur, S. P., Holt, V. L., Lee, J. H., Jiang, H. & Rust, P. F. Levels of antibodies to transferrin and alpha 2-HS glycoprotein in women with and without endometriosis. Am. J. Reprod. Immunol. 40, 69–73 (1998).

11. Lee, Y. H. et al. Limited value of pro-inflammatory oxylipins and cytokines as circulating biomarkers in endometriosis – a targeted ‘omics study. Sci. Rep. 6, 26117 (2016).

12. Mathur, S. P. Autoimmunity in endometriosis: relevance to infertility. Am. J. Reprod. Immunol. 44, 89–95 (2000).

13. Matorras, R. et al. Prevalence of endometriosis in women with systemic lupus erythematosus and Sjögren’s syndrome. Lupus 16, 736–740 (2007).

14. Jess, T., Frisch, M., Jørgensen, K. T., Pedersen, B. V. & Nielsen, N. M. Increased risk of inflammatory bowel disease in women with endometriosis: a nationwide Danish cohort study. Gut 61, 1279–1283 (2012).

15. Sinaii, N., Cleary, S. D., Ballweg, M. L., Nieman, L. K. & Stratton, P. High rates of autoimmune and endocrine disorders, fibromyalgia, chronic fatigue syndrome and atopic diseases among women with endometriosis: a survey analysis. Hum. Reprod. 17, 2715–24 (2002).

16. Matarese, G., De Placido, G., Nikas, Y. & Alviggi, C. Pathogenesis of endometriosis: natural immunity dysfunction or autoimmune disease? Trends Mol. Med. 9, 223–228 (2003).

17. Vanni, V. S. et al. Concomitant autoimmunity may be a predictor of more severe stages of endometriosis. Sci. Rep. 11, 15372 (2021).

18. Fernandéz-Shaw, S. et al. Anti-endometrial antibodies in women measured by an enzyme-linked immunosorbent assay. Hum. Reprod. 11, 1180–4 (1996).

19. el-Roeiy, A. et al. Danazol but not gonadotropin-releasing hormone agonists suppresses autoantibodies in endometriosis. Fertil. Steril. 50, 864–71 (1988).

20. Zhou, J. et al. Peritoneal Fluid Cytokines Reveal New Insights of Endometriosis Subphenotypes. Int. J. Mol. Sci. 21, 3515 (2020).

21. Hever, A. et al. Human endometriosis is associated with plasma cells and overexpression of B lymphocyte stimulator. Proc. Natl. Acad. Sci. 104, 12451–12456 (2007).

22. Ramírez-Pavez, T. N. et al. The Role of Peritoneal Macrophages in Endometriosis. Int. J. Mol. Sci. 22, 10792 (2021).

23. Jeljeli, M. et al. Macrophage Immune Memory Controls Endometriosis in Mice and Humans. Cell Rep. 33, 108325 (2020).

24. Hogg, C. et al. Macrophages inhibit and enhance endometriosis depending on their origin. Proc. Natl. Acad. Sci. 118, (2021).

25. Shi, D. & Jiang, P. A Different Facet of p53 Function: Regulation of Immunity and Inflammation During Tumor Development. Front. Cell Dev. Biol. 9, (2021).

26. Soussi, T. p53 Antibodies in the sera of patients with various types of cancer: a review. Cancer Res. 60, 1777–88 (2000).

27. Ying, T.-H. et al. Association of p53 and CDKN1A genotypes with endometriosis. Anticancer Res. 31, 4301–6 (2011).

28. Gylfason, J. T. et al. Quantitative DNA perturbations of p53 in endometriosis: analysis of American and Icelandic cases. Fertil. Steril. 84, 1388–1394 (2005).

29. Bischoff, F. Z., Heard, M. & Simpson, J. L. Somatic DNA alterations in endometriosis: high frequency of chromosome 17 and p53 loss in late-stage endometriosis. J. Reprod. Immunol. 55, 49–64 (2002).

30. Sáinz de la Cuesta, R., Izquierdo, M., Cañamero, M., Granizo, J. J. & Manzarbeitia, F. Increased prevalence of p53 overexpression from typical endometriosis to atypical endometriosis and ovarian cancer associated with endometriosis. Eur. J. Obstet. Gynecol. Reprod. Biol. 113, 87–93 (2004).

31. Komarova, E. A. et al. p53 is a suppressor of inflammatory response in mice. FASEB J. 19, 1030–1032 (2005).

32. Zheng, S.-J., Lamhamedi-Cherradi, S.-E., Wang, P., Xu, L. & Chen, Y. H. Tumor Suppressor p53 Inhibits Autoimmune Inflammation and Macrophage Function. Diabetes 54, 1423–1428 (2005).

33. Yoon, K. W. et al. Control of signaling-mediated clearance of apoptotic cells by the tumor suppressor p53. Science (80-.). 349, (2015).

34. Witalison, E., Thompson, P. & Hofseth, L. Protein Arginine Deiminases and Associated Citrullination: Physiological Functions and Diseases Associated with Dysregulation. Curr. Drug Targets 16, 700–710 (2015).

35. Rahmioglu, N. et al. World Endometriosis Research Foundation Endometriosis Phenome and Biobanking Harmonization Project: III. Fluid biospecimen collection, processing, and storage in endometriosis research. Fertil. Steril. (2014) doi:10.1016/j.fertnstert.2014.07.1208.

36. ASRM. Revised American Society for Reproductive Medicine classification of endometriosis: 1996. Fertil Steril. 67, 817–821 (1997).

37. AFS. Revised American Fertility Society classification of endometriosis: 1985. Fertil. Steril. 43, 351–352 (1985).

38. Boutell, J. M., Hart, D. J., Godber, B. L. J., Kozlowski, R. Z. & Blackburn, J. M. Functional protein microarrays for parallel characterisation of p53 mutants. Proteomics 4, 1950–1958 (2004).

39. Blackburn, J. M. & Shoko, A. Protein Function Microarrays for Customised Systems-Oriented Proteome Analysis. in 305–330 (2011). doi:10.1007/978-1-61779-286-1_21.

40. Kan, R. et al. Regulation of mouse oocyte microtubule and organelle dynamics by PADI6 and the cytoplasmic lattices. Dev. Biol. 350, 311–322 (2011).

41. Wang, S. & Wang, Y. Peptidylarginine deiminases in citrullination, gene regulation, health and pathogenesis. Biochim. Biophys. Acta - Gene Regul. Mech. 1829, 1126–1135 (2013).

42. Hensen, S. M. M. & Pruijn, G. J. M. Methods for the Detection of Peptidylarginine Deiminase (PAD) Activity and Protein Citrullination. Mol. Cell. Proteomics 13, 388–396 (2014).

43. Chen, J., Bardes, E. E., Aronow, B. J. & Jegga, A. G. ToppGene Suite for gene list enrichment analysis and candidate gene prioritization. Nucleic Acids Res. 37, W305–W311 (2009).

44. Koscielny, G. et al. Open Targets: a platform for therapeutic target identification and validation. Nucleic Acids Res. 45, D985–D994 (2017).

45. Inagaki, N. Analysis of intra-uterine cytokine concentration and matrix-metalloproteinase activity in women with recurrent failed embryo transfer. Hum. Reprod. 18, 608–615 (2003).

46. Pillai, S. et al. Antibodies to endometrial transferrin and alpha 2-Heremans Schmidt (HS) glycoprotein in patients with endometriosis. Am. J. Reprod. Immunol. 35, 483–94 (1996).

47. Gellersen, B. & Brosens, J. J. Cyclic Decidualization of the Human Endometrium in Reproductive Health and Failure. Endocr. Rev. 35, 851–905 (2014).

48. Monsivais, D. et al. Endometrial receptivity and implantation require uterine BMP signaling through an ACVR2A-SMAD1/SMAD5 axis. Nat. Commun. 12, 3386 (2021).

49. Zhang, L. et al. An immunological renal disease in transgenic mice that overexpress Fli-1, a member of the ets family of transcription factor genes. Mol. Cell. Biol. 15, 6961–6970 (1995).

50. Yokoyama, K. BANK regulates BCR-induced calcium mobilization by promoting tyrosine phosphorylation of IP3 receptor. EMBO J. 21, 83–92 (2002).

51. Kotronis, K. et al. Protein expression pattern of tissue inhibitor of metalloproteinase-3 (TIMP3) in endometriosis and normal endometrium. Gynecol. Endocrinol. 35, 1103–1106 (2019).

52. Nap, A. W., Dunselman, G. A. J., de Goeij, A. F. P. M., Evers, J. L. H. & Groothuis, P. G. Inhibiting MMP activity prevents the development of endometriosis in the chicken chorioallantoic membrane model. Hum. Reprod. 19, 2180–2187 (2004).

53. Blagih, J. et al. Cancer-Specific Loss of p53 Leads to a Modulation of Myeloid and T Cell Responses. Cell Rep. 30, 481-496.e6 (2020).

54. Cooks, T., Harris, C. C. & Oren, M. Caught in the cross fire: p53 in inflammation. Carcinogenesis 35, 1680–1690 (2014).

55. Ciesielski, O. et al. Citrullination in the pathology of inflammatory and autoimmune disorders: recent advances and future perspectives. Cell. Mol. Life Sci. 79, 94 (2022).

56. Carp, H. J. A., Selmi, C. & Shoenfeld, Y. The autoimmune bases of infertility and pregnancy loss. J. Autoimmun. 38, J266–J274 (2012).

57. Lang, G. A. & Yeaman, G. R. Autoantibodies in Endometriosis Sera Recognize a Thomsen–Friedenreich-like Carbohydrate Antigen. J. Autoimmun. 16, 151–161 (2001).

58. Dmowski, W. P. et al. The effect of endometriosis, its stage and activity, and of autoantibodies on in vitro fertilization and embryo transfer success rates**Presented at the Conjoint Meeting of The American Fertility Society and The Canadian Fertility and Andrology Society, M. Fertil. Steril. 63, 555–562 (1995).

59. Gajbhiye, R. et al. Multiple endometrial antigens are targeted in autoimmune endometriosis. Reprod. Biomed. Online 16, 817–824 (2008).

60. Kennedy, S. H., Sargent, I. L., Starkey, P. M., Hicks, B. R. & Barlow, D. H. Localization of anti-endometrial antibody binding in women with endometriosis using a double-labelling immunohistochemical method. Br. J. Obstet. Gynaecol. 97, 671–4 (1990).

61. Badaway, S. Z., Cuenca, V., Freliech, H. & Stefanu, C. Endometrial antibodies in serum and peritoneal fluid of infertile patients with and without endometriosis. Fertil. Steril. 53, 930–2 (1990).

62. Taylor, P. V et al. Autoreactivity in women with endometriosis. Br. J. Obstet. Gynaecol. 98, 680–4 (1991).

63. Wild, R. A. & Shivers, C. A. Antiendometrial antibodies in patients with endometriosis. Am. J. Reprod. Immunol. Microbiol. 8, 84–6 (1985).

64. Pauklin, S., Sernández, I. V., Bachmann, G., Ramiro, A. R. & Petersen-Mahrt, S. K. Estrogen directly activates AID transcription and function. J. Exp. Med. 206, 99–111 (2009).

65. Nagata, S., Hanayama, R. & Kawane, K. Autoimmunity and the Clearance of Dead Cells. Cell 140, 619–630 (2010).

66. Elliott, M. R. & Ravichandran, K. S. Clearance of apoptotic cells: implications in health and disease. J. Cell Biol. 189, 1059–1070 (2010).

67. Lee, Y. H. et al. Dysregulated sphingolipid metabolism in endometriosis. J. Clin. Endocrinol. Metab. 99, E1913–21 (2014).

68. Weedin, E. et al. Gonadotropin-releasing hormone receptor autoantibody activity in polycystic ovary syndrome - stability of autoantibody levels over time. Fertil. Steril. 110, e113 (2018).

69. Li, Y. et al. Longitudinal serum autoantibody repertoire profiling identifies surgery-associated biomarkers in lung adenocarcinoma. EBioMedicine 53, 102674 (2020).

70. Lu, H., Goodell, V. & Disis, M. L. Humoral Immunity Directed against Tumor-Associated Antigens As Potential Biomarkers for the Early Diagnosis of Cancer. J. Proteome Res. 7, 1388–1394 (2008).

71. Anderson, K. S. et al. Autoantibody Signature for the Serologic Detection of Ovarian Cancer. J. Proteome Res. 14, 578–586 (2015).

72. von Stemann, J. H. et al. Cytokine autoantibodies are stable throughout the haematopoietic stem cell transplantation course and are associated with distinct biomarker and blood cell profiles. Sci. Rep. 11, 23971 (2021).

73. Tan, Y. et al. Single cell analysis of endometriosis reveals a coordinated transcriptional program driving immunotolerance and angiogenesis across eutopic and ectopic tissues. BioRivx doi:10.1101/2021.07.28.453839.

74. Olkowska-Truchanowicz, J. et al. Endometriotic Peritoneal Fluid Stimulates Recruitment of CD4+CD25highFOXP3+ Treg Cells. J. Clin. Med. 10, 3789 (2021).

75. Guo, M. et al. Mass cytometry analysis reveals a distinct immune environment in peritoneal fluid in endometriosis: a characterisation study. BMC Med. 18, 3 (2020).

76. Olkowska-Truchanowicz, J. et al. CD4+ CD25+ FOXP3+ regulatory T cells in peripheral blood and peritoneal fluid of patients with endometriosis. Hum. Reprod. 28, 119–124 (2013).

77. Podgaec, S., Rizzo, L. V., Fernandes, L. F. C., Baracat, E. C. & Abrao, M. S. CD4+CD25highFoxp3+ Cells Increased in the Peritoneal Fluid of Patients with Endometriosis. Am. J. Reprod. Immunol. 68, 301–308 (2012).

78. Annibali, V. et al. CD161highCD8+T cells bear pathogenetic potential in multiple sclerosis. Brain 134, 542–554 (2011).

79. Uhlén, M. et al. Tissue-based map of the human proteome. Science (80-.). 347, (2015).

80. Protein Altas. https://www.proteinatlas.org/ENSG00000141510-TP53/tissue/endometrium.

81. Dufournet, C. et al. Expression of apoptosis-related proteins in peritoneal, ovarian and colorectal endometriosis. J. Reprod. Immunol. 70, 151–162 (2006).

82. Laudanski, P. et al. Expression of selected tumor suppressor and oncogenes in endometrium of women with endometriosis. Hum. Reprod. 24, 1880–1890 (2009).

83. Goumenou, A. et al. Immunohistochemical Expression of p53, MDM2, and p21Wafi Oncoproteins in Endometriomas But Not Adenomyosis. J. Soc. Gynecol. Investig. 12, 263–266 (2005).

84. Katchman, B. A. et al. Proteomic mapping of p53 immunogenicity in pancreatic, ovarian, and breast cancers. PROTEOMICS - Clin. Appl. 10, 720–731 (2016).

85. Leroy, B., Anderson, M. & Soussi, T. TP53 Mutations in Human Cancer: Database Reassessment and Prospects for the Next Decade. Hum. Mutat. 35, 672–688 (2014).

86. Herkel, J. et al. Autoimmunity to the p53 Protein is a Feature of Systemic Lupus Erythematosus (SLE) Related to Anti-DNA Antibodies. J. Autoimmun. 17, 63–69 (2001).

87. Tsou, C.-L. et al. Critical roles for CCR2 and MCP-3 in monocyte mobilization from bone marrow and recruitment to inflammatory sites. J. Clin. Invest. 117, 902–909 (2007).

88. Lee, C.-Y. et al. Mining the Human Tissue Proteome for Protein Citrullination. Mol. Cell. Proteomics 17, 1378–1391 (2018).

89. Tilvawala, R. et al. The Rheumatoid Arthritis-Associated Citrullinome. Cell Chem. Biol. 25, 691-704.e6 (2018).

90. Guerrin, M. et al. cDNA cloning, gene organization and expression analysis of human peptidylarginine deiminase type I. Biochem. J. 370, 167–174 (2003).

91. Chavanas, S. et al. Comparative analysis of the mouse and human peptidylarginine deiminase gene clusters reveals highly conserved non-coding segments and a new human gene, PADI6. Gene 330, 19–27 (2004).

92. O’Donoghue, K. Identification of fetal mesenchymal stem cells in maternal blood: implications for non-invasive prenatal diagnosis. Mol. Hum. Reprod. 9, 497–502 (2003).

93. Evans, P. C. et al. Long-term fetal microchimerism in peripheral blood mononuclear cell subsets in healthy women and women with scleroderma. Blood 93, 2033–7 (1999).

94. Ando, T., Imaizumi, M., Graves, P. N., Unger, P. & Davies, T. F. Intrathyroidal Fetal Microchimerism in Graves’ Disease. J. Clin. Endocrinol. Metab. 87, 3315–3320 (2002).

95. Nnoaham, K. E. et al. Impact of endometriosis on quality of life and work productivity: a multicenter study across ten countries. Fertil. Steril. 96, 366-373.e8 (2011).

96. Lee, Y. H. et al. Elevated peritoneal fluid ceramides in human endometriosis-associated infertility and their effects on mouse oocyte maturation. Fertil. Steril. 110, 767-777.e5 (2018).

97. Yeung, H.-Y. & Dendrou, C. A. Pregnancy Immunogenetics and Genomics: Implications for Pregnancy-Related Complications and Autoimmune Disease. Annu. Rev. Genomics Hum. Genet. 20, 73–97 (2019).

98. Janssen, E., Zhu, M., Zhang, W., Koonpaew, S. & Zhang, W. LAB: A new membrane-associated adaptor molecule in B cell activation. Nat. Immunol. 4, 117–123 (2003).

99. Latifi, Z. et al. Dual role of TGF-β in early pregnancy: clues from tumor progression. Biol. Reprod. 100, 1417–1430 (2019).

100. Young, V. J., Ahmad, S. F., Duncan, W. C. & Horne, A. W. The role of TGF-β in the pathophysiology of peritoneal endometriosis. Hum. Reprod. Update 23, 548–559 (2017).

101. Germeyer, A., Savaris, R. F., Jauckus, J. & Lessey, B. Endometrial beta3 Integrin profile reflects endometrial receptivity defects in women with unexplained recurrent pregnancy loss. Reprod. Biol. Endocrinol. 12, 53 (2014).

