## Supplementary material for "Peritoneal autoantibody landscape in endometriosis": Figure S1

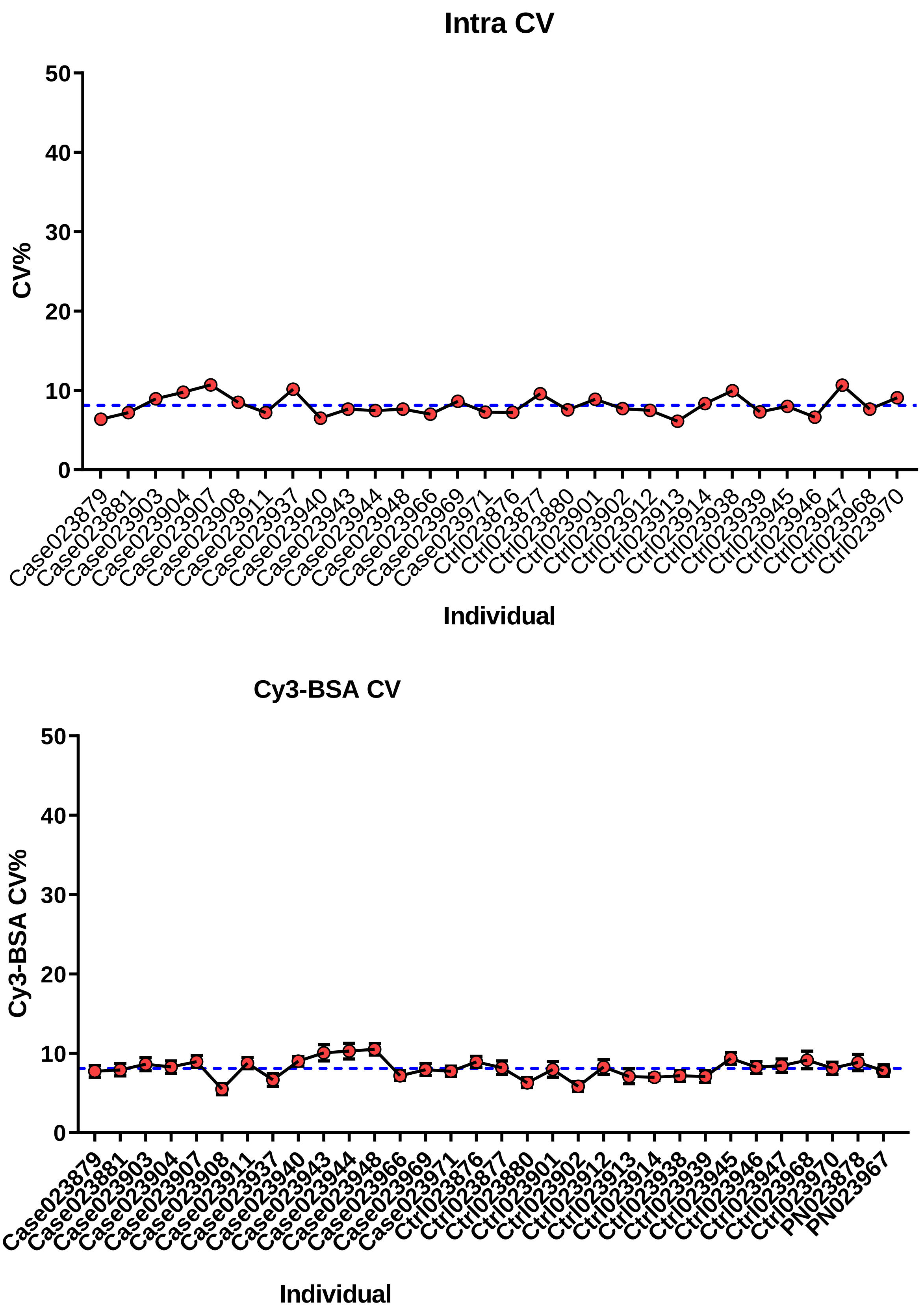


**Figure S1. Robust quantitation of autoantibody protein array.** (A) Mean of CV% of all protein replica spots across all samples was 8.12%. (B) Cy3-labelled biotinylated BSA (Cy3-BSA), serves as the positive control across slides and as a housekeeping probe for normalisation of signal intensities across samples, has a CV% of 8.08%.
