## Supplementary figures and images for "Peritoneal autoantibody landscape in endometriosis"

### Figure S2

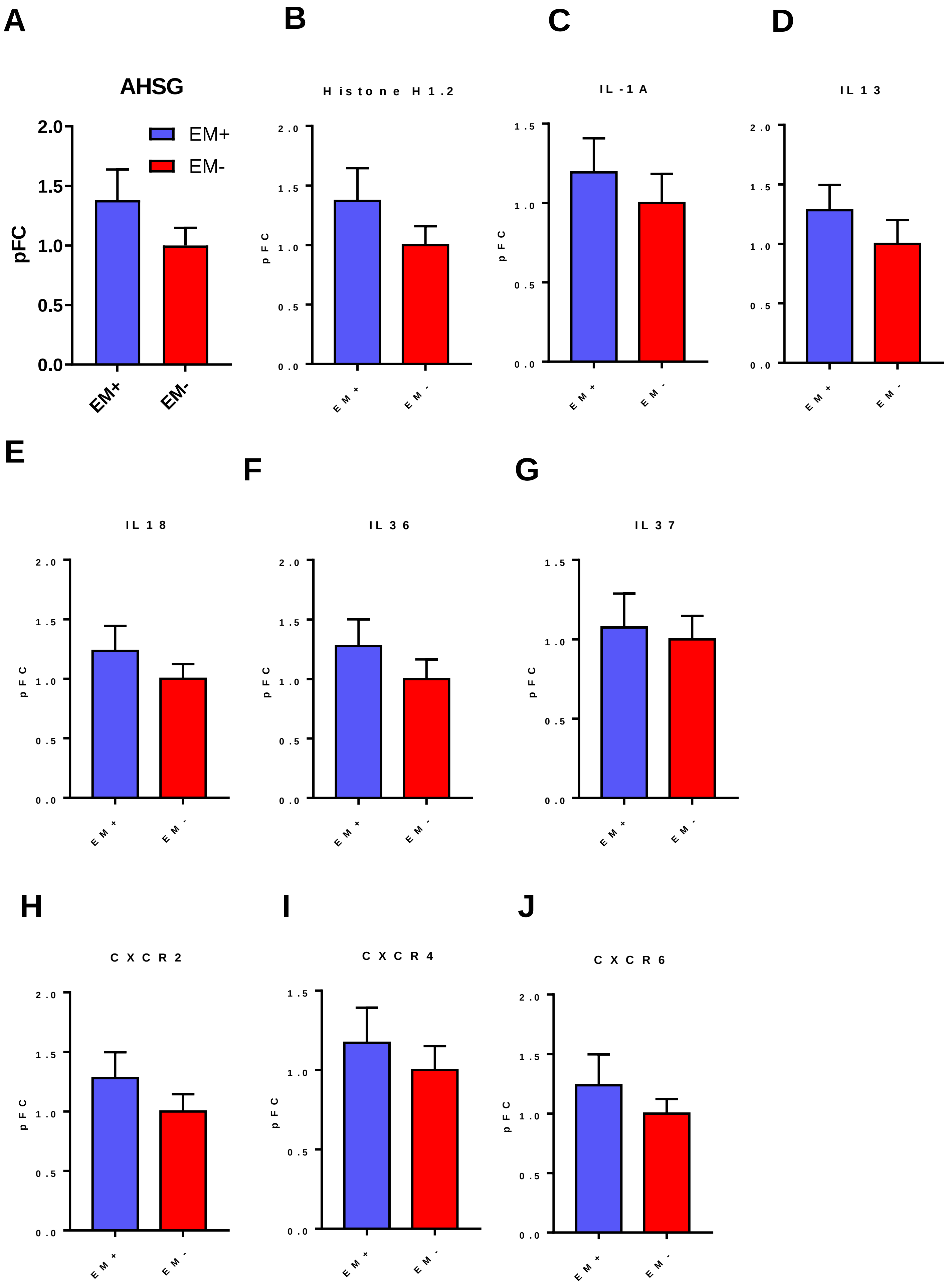


**Figure S2. Bar graphs of peritoneal fluid cytokine autoantibody levels in endometriosis.**
