## Supplementary material for "Peritoneal autoantibody landscape in endometriosis": Figure S3

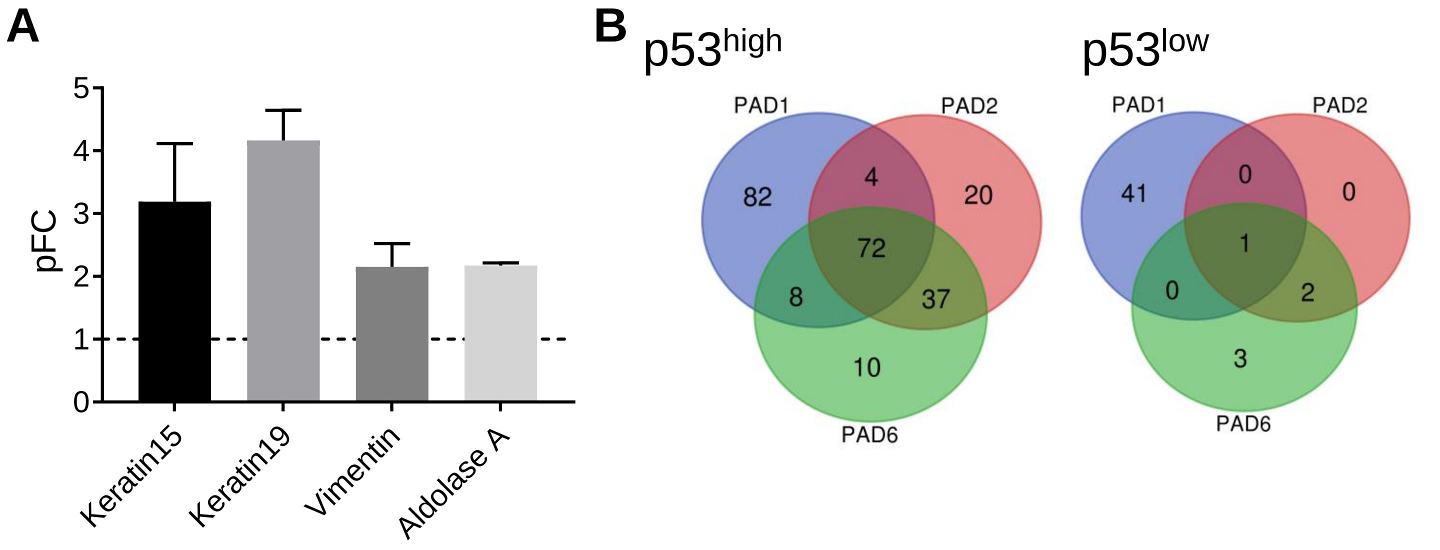


**Figure S3. Validation of citrullinated autoantibody protein array.** (A) Known citrullinated proteins keratins (KRT15 and KRT19), vimentin (VIM) and aldolase (ALDOA) were detected using our developed citrullinated autoantibody protein array which validated the assay. (B) The anti-p53 autoantibody groups had different citrullination patterns depending on whether they were incubated with PAD1, 2, or 6, with PAD1 generating the most reactive autoantigens. In the $p53^{high}$ group 72 citrullinated autoantibodies overlapped in autoantibodies generated by the three PADs and in the $p53^{low}$ group only one overlapped.
